# Fish predation on corals promotes the dispersal of coral symbionts

**DOI:** 10.1101/2020.08.10.243857

**Authors:** Carsten G.B. Grupstra, Kristen M. Rabbitt, Lauren I. Howe-Kerr, Adrienne M.S. Correa

## Abstract

Predators drive top-down effects that shape prey communities, but the role of predators in dispersing prey microbiomes is rarely examined. We tested whether coral-eating (corallivorous) fish disperse the single-celled dinoflagellate symbionts (family Symbiodiniaceae) of their prey. Our findings demonstrate that: (1) coral-eating fish egest feces containing live Symbiodiniaceae at densities up to seven orders of magnitude higher than other environmental reservoirs such as sediments and water; (2) Symbiodiniaceae communities in the feces of most corallivores are compositionally similar to those in corals; (3) some obligate corallivore species release over 100 million Symbiodiniaceae cells per 100 m^2^ per day; and (4) after being egested, corallivore feces often come in direct contact with coral colonies (potential hosts for Symbiodiniaceae). These findings suggest that fish predators can play an important role in symbiont acquisition by corals; such predators may have a previously unrecognized, indirect positive effect on prey health.

## Introduction

All animals and plants host microbiomes - resident communities of microorganisms – that influence their health and survival (Bosch and McFall-Ngai, 2011; McFall-Ngai et al., 2013). Many hosts establish or modify their microbiome by taking up microorganisms from the environment (Moran and Sloan, 2015). Stony corals, for example, harbor single-celled dinoflagellates in the family Symbiodiniaceae and utilize their photosynthetic products to fuel the construction of reef frameworks (Muscatine et al., 1984). Approximately 75% of spawning and 10% of brooding coral species acquire their Symbiodiniaceae from the environment with each generation (Baird et al., 2009), and adult corals may take up environmental Symbiodiniaceae cells to replenish their microbiomes following abiotic stress (Boulotte et al., 2016; Lewis and Coffroth, 2004; Rouzé et al., 2019). However, the dispersal mechanisms that make Symbiodiniaceae cells available to prospective host corals have not been resolved (but see Coffroth et al., 2006; Nitschke et al., 2016), and Symbiodiniaceae cell densities in environmental reservoirs appear relatively low (sediments: 10^1^-10^3^ cells ml^−1^; seawater: 10^0^-10^1^ cells ml^−1^; macroalgae: 10^2^-10^3^ cells ml^−1^, Figure 1; Castro-Sanguino and Sánchez, 2012; Fujise et al., *in review*; Littman et al., 2008).

**Figure 1.**
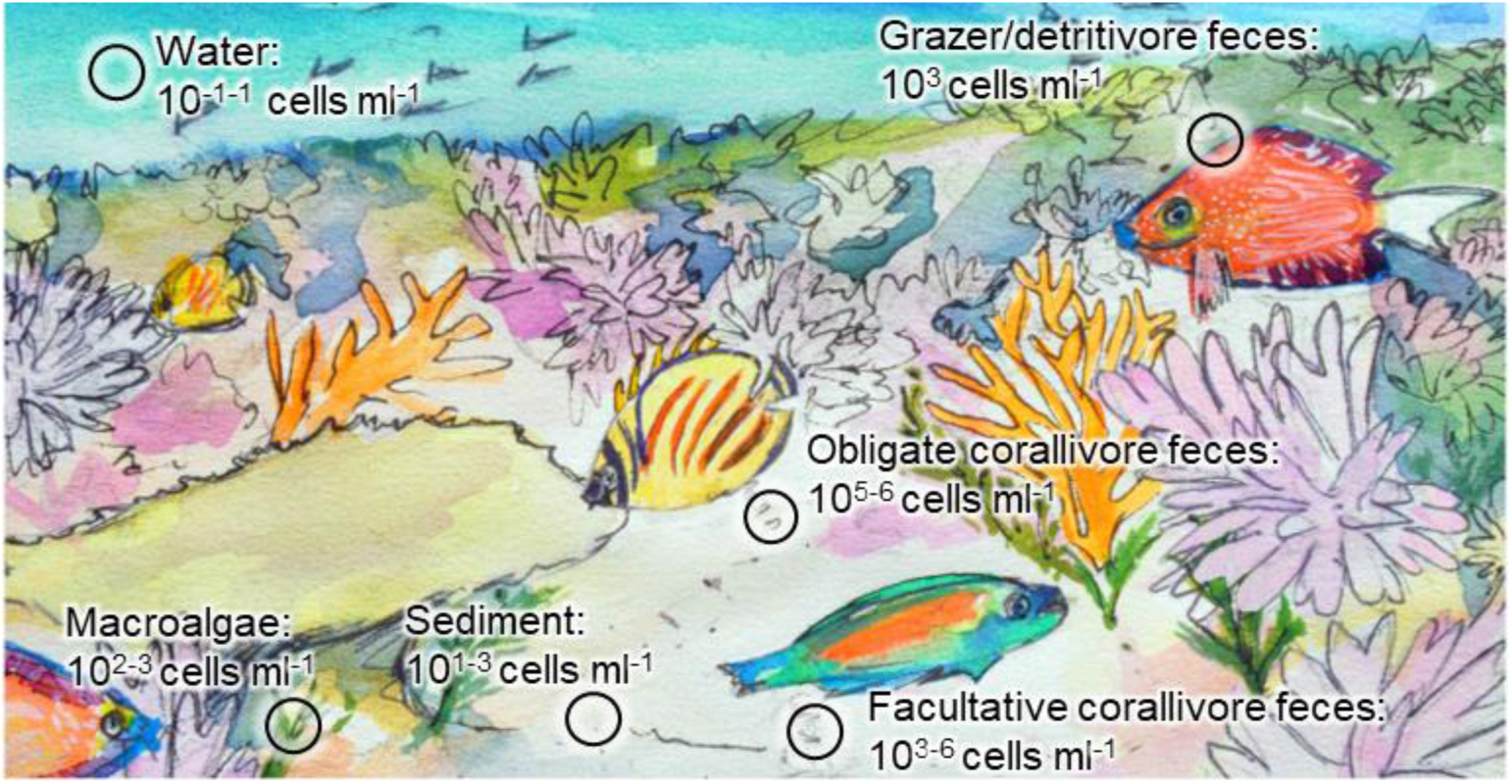
Generalized mean Symbiodiniaceae cell densities in coral reef environmental reservoirs. Cell densities reported in this study (all obligate corallivore and grazer/detritivore feces data and some data for facultative corallivore feces, sediments and seawater) represent live cell densities only, based on results from hemocytometry with the cell viability dye, trypan blue. Previously published cell densities for sediment and seawater (Fujise et al., *in review*; Littman et al., 2008), macroalgae (Fujise et al., *in review*), and facultative corallivore feces (Castro-Sanguino and Sánchez, 2012) may include both dead and live cells and were quantified using hemocytometry (Castro-Sanguino and Sánchez, 2012), a combination of flow-cytometry and hemocytometry (Littman et al., 2008), or quantitative PCR (Fujise et al., *in review*).

Consumers (e.g., nectarivores, herbivores, predators) are known to mediate the dispersal of microbiome constituents for various host organisms. Nectar-associated yeasts and bacteria, for example, are transmitted by nectarivores (Brysch-Herzberg, 2004), and fungi associated with plant mycorrhiza are dispersed in the feces of diverse herbivorous animals (Vašutová et al., 2019). Corallivores (coral-eating fish and invertebrates) that feed on corals without killing them may similarly mediate the dispersal of coral microbiomes by egesting Symbiodiniaceae - and other members of the coral microbiome - in their feces as they move across reefs (Muller Parker, 1984; Smriga et al., 2010). A total of 128 fish species spanning 11 families consume corals as part (i.e., facultative corallivores) or all (i.e., obligate corallivores) of their diets, and most of them occur in the Indo-Pacific or the Red Sea (Cole et al., 2008; Rotjan and Lewis, 2008). To date, dispersal of Symbiodiniaceae has been quantified from one fish species: the Caribbean facultative corallivorous parrotfish *Sparisoma viride* (Castro-Sanguino and Sánchez, 2012). Symbiodiniaceae cells were present at low abundances (similar to sediments; Castro-Sanguino and Sánchez, 2012; Littman et al., 2008) in analyzed feces, and a Symbiodiniaceae genus (i.e*., Cladocopium*) that was locally dominant in corals was not detected (Castro-Sanguino and Sánchez, 2012). These results suggested that facultative corallivores are not major dispersers of Symbiodiniaceae. However, obligate corallivores ingest higher abundances of coral tissue than facultative corallivores and may therefore be more important drivers of Symbiodiniaceae dispersal, but this has not been previously studied. We conducted the first comprehensive, ecological-scale quantification of fish-mediated coral microbiome dispersal on a Pacific reef and found, surprisingly, that the feces of *both* obligate and facultative corallivorous fishes contain high densities of live Symbiodiniaceae cells. Thus, this study adds a new dimension to our understanding of the potential outcomes of predation and reveals an important dispersal mechanism for Symbiodiniaceae on Pacific reefs.

We characterized Symbiodiniaceae densities and community compositions in the feces of four obligate corallivores (the butterflyfishes *Chaetodon lunulatus*, *Chaetodon ornatissimus*, *Chaetodon reticulatus*, and the filefish *Amanses scopas)* and three facultative corallivores (the butterflyfishes *Chaetodon citrinellus* and *Chaetodon pelewensis*, and the parrotfish *Chlorurus spilurus*) from a reefscape in Mo’orea, French Polynesia (Cole et al., 2008; Ezzat et al., 2020; Rotjan and Lewis, 2008). To test whether the feces of obligate corallivores constitute ‘hotspots’ of live Symbiodiniaceae and are proximate environmental sources of Symbiodiniaceae for prospective coral host colonies, we additionally characterized Symbiodiniaceae from reef-associated sediments and water, as well as the feces of two grazer/detritivores (i.e., surgeonfishes *Ctenochaetus flavicauda* and *Ctenochaetus striatus*). We then compared Symbiodiniaceae communities in all of these samples to those harbored by the locally dominant reef-building corals *Acropora hyacinthus*, *Pocillopora* species complex (Forsman et al., 2015, 2009) and *Porites lobata* species complex (Gélin et al., 2017). Next, we used bootstrap models based on *in situ* surveys and observations to generate the first reef-scale predictions of Symbiodiniaceae dispersal for three corallivorous fish species. Finally, we quantified how often egested feces came in direct contact with coral colonies *in situ* to estimate how frequently fish predation and egestion potentially influences Symbiodiniaceae assembly in coral hosts.

## Results and Discussion

Using a cell viability dye (Figure S1; Strychar and Coates, 2004; Zhang et al., 2008), we detected live Symbiodiniaceae cells in all obligate corallivore feces (n=40, across four species) and in ~81% of facultative corallivore feces (n=18 of 22, across three species). Live Symbiodiniaceae cells were detected in ~36% of grazer/detritivore feces (n=5 of 14, across two species), ~42% of sediment samples (n=5 of 12) and ~8% of seawater samples (n=1 of 12). The long-term viability of Symbiodiniaceae cells in feces was further confirmed through culturing (attempted from 61 fish fecal pellets using 24 replicate wells per sample; 1464 wells total). Six to ten weeks after cultures were instigated, live Symbiodiniaceae cells in coccoid, mitotic and motile life stages were documented in at least one replicate culture well from 54% of obligate corallivore feces (n=19 of 35 samples), 11% of facultative corallivore feces (n=2 of 18 samples), and 38% of detritivore feces (n=3 of 8 samples) indicating that consumer feces contain Symbiodiniaceae cells that are competent for long-term survival and division.

Live Symbiodiniaceae cell densities differed among obligate corallivore feces, facultative corallivore feces, and detritivore feces, reef sediments and seawater (overall Kruskal-Wallis test results: Chi-squared=74.3, df=3, p-value<0.001, Figure 2). The mean (± 95% CI) cell concentration ml^−1^ of feces was 5.30⋅10^6^ ± 1.29⋅10^6^ for obligate corallivores; this was ~5.6 times higher than in facultative corallivore feces (9.46⋅10^5^ ± 7.51⋅10^5^; pairwise test statistics in Table S2, S3), ~750 times higher than in grazer/detritivore feces (7.08⋅10^3^ ± 6.65⋅10^3^) and ~460,000 times higher than in sediment and seawater samples combined (11.5 ± 5.50). Mean live Symbiodiniaceae cell concentrations in feces of individual obligate corallivore species (Figure 2-figure supplement 1) ranged from 7.96⋅10^5^ ± 5.1⋅10^5^ (*Amanses scopas*) to 7.6⋅10^6^ ± 2.5⋅10^6^ (*Chaetodon reticulatus*) ml^−1^ of feces and were 4-5 orders of magnitude higher than sediment samples (22.75 ± 22.37) and 6-7 orders of magnitude higher than seawater samples (0.23 ± 0.51). Mean live Symbiodiniaceae cell densities in feces of individual facultative corallivore species were more variable than obligate corallivores (Figure 2-figure supplement 1) and ranged from 2.1⋅10^4^ ± 2.8⋅10^4^ (*Chlorurus spilurus*) to 2.1⋅10^6^ ± 1.9⋅10^6^ (*Chaetodon pelewensis*) ml^−1^ feces. Overall, live Symbiodiniaceae cell densities in corallivore feces in this study were 1-3 orders of magnitude higher than values previously reported from the corallivorous parrotfish *Sparisoma viride* (live and dead cells combined: 3.2⋅10^3^-8.9⋅10^3^ cells ml^−1^; Castro-Sanguino and Sánchez, 2012), even for the most comparable species, *Chlorurus spilurus*, which is also a facultatively corallivorous parrotfish. These higher cell densities are likely due to dietary differences among fishes. The corallivores examined in our study generally rely on corals for a larger part of their diets (Harmelin-Vivien and Bouchon-Navaro, 1983) than *S. viride* (but note that no data are available for C. spilurus; Castro-Sanguino and Sánchez, 2012), and therefore ingest higher abundances of Symbiodiniaceae cells.

**Figure 2.**
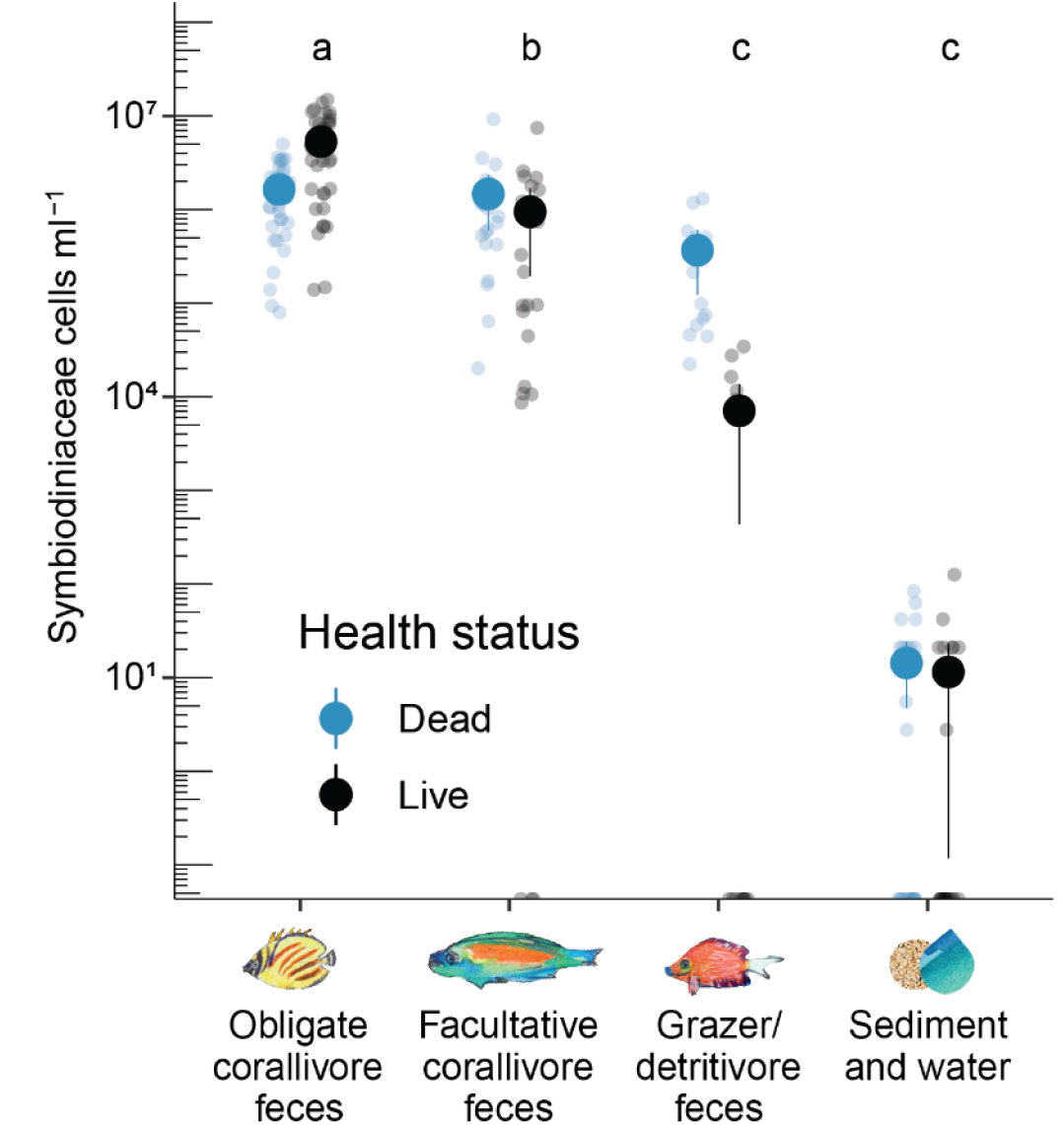
Corallivore (coral-eating animal) feces contain live Symbiodiniaceae cells at densities several orders of magnitude higher than in grazer/detritivore (detritus-eating animal) feces and reef-associated sediment and seawater. Large dots indicate the mean densities of dead (blue) and live (black) cells for each sample category; vertical lines depict 95% confidence intervals. Small dots indicate dead (blue) and live (black) cell densities from individual samples. Letters indicate significant differences (p<0.05) in live Symbiodiniaceae cell densities based on pairwise Dunn tests with Bonferroni-corrected p-values (overall Kruskal-Wallis test results: Chi-squared=74.3, df=3, p-value<0.001; see Table S2 for pairwise test results). Included species (and sample sizes) for each sample category are as follows: Obligate corallivores: *Amanses scopas*, AMSC (7)*; Chaetodon lunulatus*, CHLU (8); *Chaetodon ornatissimus*, CHOR (14); *Chaetodon reticulatus*, CHRE (11). Facultative corallivores: *Chaetodon pelewensis*, CHPE (8); *Chaetodon citrinellus*, CHCI (6); and *Chlorurus spilurus* CHSP (8). Grazer/Detritivores: *Ctenochaetus flavicauda*, CTFL (8); and *Ctenochaetus striatus*, CTST (6). Sediment and water: Sediment, SED (12); Water, WAT (12). See Table S1 for additional information on replication. For data used see **Figure 2-Source Data 1**.

Amplicon sequencing of the Symbiodiniaceae internal transcribed spacer-2 (ITS-2) region of rDNA demonstrated that, at the genus level, symbiont communities differed among coral species, obligate corallivore feces, grazer/detritivore feces, and facultative corallivore feces, reef sediments and seawater (PERMANOVA: df=6, F=17.3, R^2^=0.58, p=0.001; Figure 3). Symbiodiniaceae communities in obligate corallivore feces overall most closely resembled those in the *Pocillopora* species complex and *Porites lobata* species complex corals (Table S4 and S5). On average, sequenced reads from obligate corallivore feces were dominated by similarities to the genus *Cladocopium* (72-98% of reads), with low relative abundances of reads identified as the genus *Durusdinium* (2-27% of reads) and *Symbiodinium* (0-1% of reads). These findings are consistent with observations (this study and Pratchett, 2014) that corallivores at our study sites mainly feed on pocilloporid and poritid corals. Facultative corallivore feces were similar to sediment and seawater samples; these feces contained, on average, less *Cladocopium* (26-81%), but more *Durusdinium* (11-73%) and *Symbiodinium* (0-8%) than obligate corallivores (Figure 3). Grazer/detritivore feces were distinct from all other sample categories. The Symbiodiniaceae genera *Breviolum* and *Fugacium*, which are rare in Mo’orean corals (Putnam et al., 2012), were detected only in grazer/detritivore feces (*Fugacium* only), reef-associated seawater (*Breviolum* only) and sediment samples.

**Figure 3.**
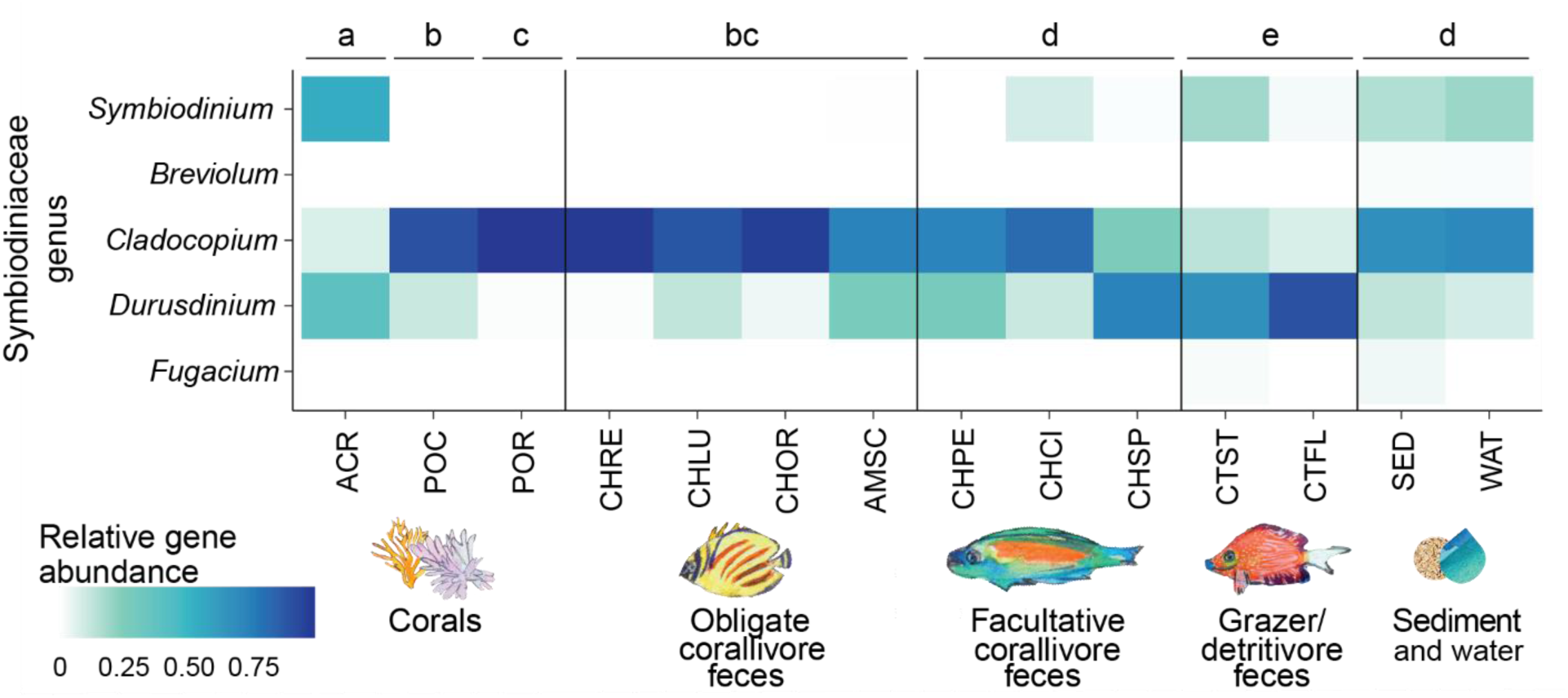
The communities of Symbiodiniaceae in obligate corallivore (coral-eating animal) feces are most similar to the communities of Symbiodiniaceae in two locally abundant coral species. Relative abundances of Internal Transcribed Spacer-2 (ITS-2) region rDNA gene variants from five Symbiodiniaceae genera in corals, obligate corallivore feces, facultative corallivore feces, grazer/detritivore feces, and reef-associated sediment and water (all included samples had >1,000 reads). Different letters indicate significant differences in Symbiodiniaceae community compositions based on pairwise PERMANOVA tests (p<0.05, using the Benjamini-Hochberg correction for multiple comparisons) using Bray-Curtis distances based on randomly subsampled (n=12) untransformed data (Table S1, S5). Overall PERMANOVA test results: df=6, F=17.3, R^2^=0.58, p=0.001. Included samples (and sample sizes) for each sample category in the figure are as follows: Corals: *Acropora hyacinthus*, ACR (11); *Pocillopora* species complex, POC (12); *Porites lobata* species complex, POR (12). Obligate corallivores: *Amanses scopas*, AMSC (7)*; Chaetodon lunulatus*, CHLU (8); *Chaetodon ornatissimus*, CHOR (14); *Chaetodon reticulatus*, CHRE (11). Facultative corallivores: *Chaetodon pelewensis*, CHPE (8); *Chaetodon citrinellus*, CHCI (6); and *Chlorurus spilurus* CHSP (8). Grazer/Detritivores: *Ctenochaetus flavicauda*, CTFL (6); and *Ctenochaetus striatus*, CTST (6). Sediment and water: Sediment, SED (12); Water, WAT (7). For data used see **Figure 3-Source Data 1**.

To generate the first estimates of the daily dispersal of live Symbiodiniaceae cells by corallivorous fish at the reef scale, we applied a bootstrap approach to *in situ* field observations (egestion rates, fecal pellet sizes), *ex situ* measurements (live Symbiodiniaceae cell densities, fecal pellet densities), and MCR LTER (Mo’orea Coral Reef Long Term Ecological Research) time series data (fish densities) for two obligate corallivores (*C. ornatissimus* and *C. reticulatus*) and one facultative corallivore (*C. citrinellus*). The mean (± 95% CI) estimated dispersal rates for the obligate corallivores were 1.01⋅10^8^ ± 6.5⋅10^6^ and 1.27⋅10^8^ ± 4.61⋅10^6^ cells per 100 m^2^ d^−1^, respectively (Figure 4); these were three orders of magnitude higher than the estimated mean for the facultative corallivore (3.32⋅10^5^ ± 6.21⋅10^4^). Differences between obligate versus facultative corallivore estimates were mostly driven by higher densities of *C. ornatissimus* and *C. reticulatus* individuals at our study site (4.7-8.8 times higher than *C. citrinellus*, Table S6), and higher Symbiodiniaceae densities in their feces (3.7-7.5 times higher than *C. citrinellus*, Table S6, Figure 2-figure supplement 1).

**Figure 4.**
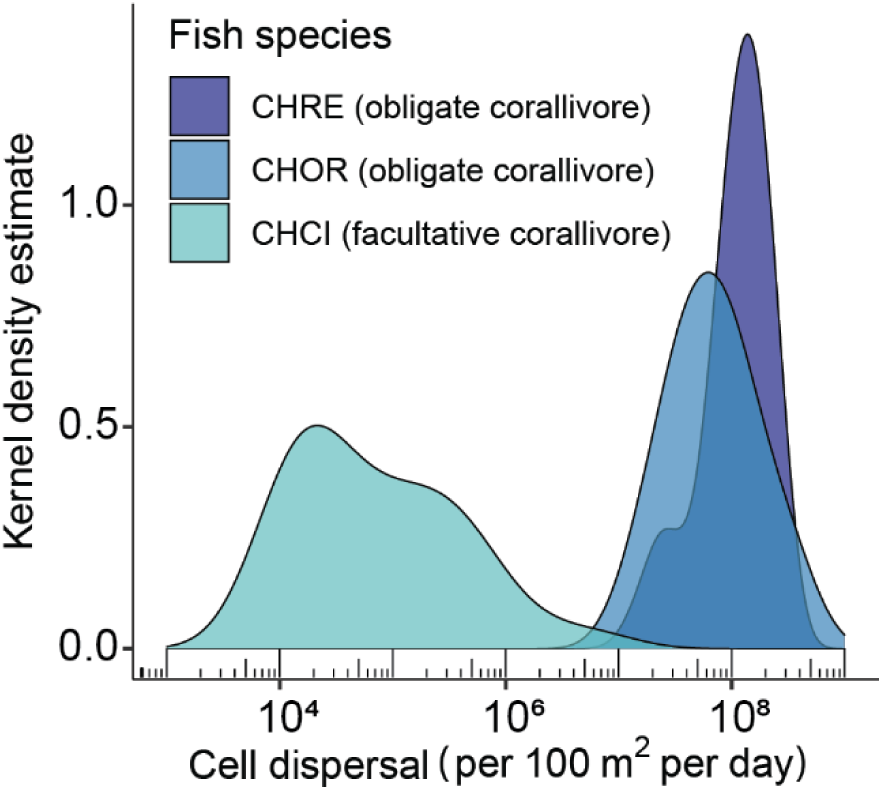
Corallivores on a South Pacific reef disperse millions of live Symbiodiniaceae cells per 100 m^2^ per day. We estimated the number of live Symbiodiniaceae cells dispersed by the obligate corallivores *Chaetodon ornatissimus* (CHOR) and *Chaetodon reticulatus* (CHRE) and the facultative corallivore *Ctenochaetus citrinellus* (CHCI) per 100 m^2^ per day by applying a bootstrap approach (1000 iterations) to the equation *T* = *eSWCF*. The estimated number of dispersed Symbiodiniaceae cells (*T*) is the product of five variables: A fish species-specific constant representing the estimated number of egestions in an eight-hour day (*e*); observed fecal pellet sizes in cm (*S*); measured densities of fecal samples in g cm^−1^ (*W*); measured densities of Symbiodiniaceae cells g^−1^ feces (*C*); and observed fish densities per 100 m^2^ (*F*). Due to variation in fish species distribution, most data for CHOR and CHRE were collected on the fore reef, whereas data for CHCI were collected on the back reef. See Table S6 for replication. For data used see **Figure 4-Source Data 1**.

This work demonstrates that obligate corallivore feces contain live Symbiodiniaceae cells at densities two to seven orders of magnitude higher than estimates for other environmental reservoirs such as the water column, sediments and macroalgae (this work; Fujise et al., *in review*; Littman et al., 2008), and up to three orders of magnitude higher than feces of the one other corallivorous fish that has been examined: the Caribbean parrotfish *S. viride* (Castro-Sanguino and Sánchez, 2012). During our surveys, we observed that egested feces often fell apart into several segments, and segments of 91% of egested feces fell onto live corals at our fore reef site (n=20 of 22 feces). Segments of only 10% of feces landed on live corals at our back reef site (n=3 of 30 feces), which is consistent with lower mean (± SD) live coral cover in this reef zone (44.8% ± 10.8 coral cover on the fore reef; 15.3% ± 7.10 back reef, ANOVA, df=1, F=44.91, p<0.001). These results indicate that fish predators commonly mediate the dispersal of Symbiodiniaceae cells to prospective coral host colonies on Pacific reefs, especially in areas with relatively high coral cover.

Our findings suggest that the feces of obligate corallivores and at least some facultative corallivores constitute significant but underexplored environmental ‘hotspots’ of Symbiodiniaceae on coral reefs; such feces can supply Symbiodiniaceae cells to potential hosts directly, or to other environmental reservoirs as they disintegrate (Castro-Sanguino and Sánchez, 2012; Nitschke et al., 2016). Corals have been shown to take up Symbiodiniaceae from sediments and seawater (Coffroth et al., 2006; Lewis and Coffroth, 2004; Nitschke et al., 2016). Thus, corals are likely capable of incorporating Symbiodiniaceae from fecal material as well. Although experimental validation for stony corals is needed, this has already been demonstrated for the sea anemone *Aiptasia pulchella* (Muller Parker, 1984). Taken together, our findings broadly suggest that some predators have indirect positive effects on prey health; such predators supply prey with microorganisms that support their function and, ultimately, survival.

The dispersal of beneficial microorganisms may become increasingly important (Foo et al., 2017; Rebollar et al., 2016) as anthropogenic stressors disrupt animal and plant microbiomes, leading to disease and mortality (Hughes et al., 2017; Jiménez and Sommer, 2017; Petton et al., 2015; Wilkinson, 2008). Our results suggest that shifts and declines in fish communities due to overfishing (Bellwood and Choat, 2011; Hawkins et al., 2007; McClanahan et al., 1999) and habitat degradation (Pratchett et al., 2006; Viviani et al., 2019) may contribute to an unexplored issue on reefs: reduced dispersal of Symbiodiniaceae, a key member of the coral microbiome. Coral-Symbiodiniaceae partnerships have been increasingly disrupted over the past four decades, resulting in coral reef decline (Hughes et al., 2017; Wilkinson, 2008). To help corals tolerate stress and mitigate reef degradation (Peixoto et al., 2017), probiotic solutions of beneficial Symbiodiniaceae (Morgans et al., 2020) and bacteria (Rosado et al., 2019) are currently being developed. We show that corallivores egest feces containing high densities of live Symbiodiniaceae cells directly onto corals. This behavior may constitute a routine, global-scale ‘restoration effort’ that inoculates corals with natural probiotics derived from nearby colonies. It is essential that healthy fish assemblages are protected and maintained on reefs as we investigate the potentially stabilizing effect of corallivores on coral microbiomes, and by extension, coral cover on reefs.

## Methods

Individuals of nine fish species, as well as corals, sediments, and seawater were collected (n=6-14 per sample type or species, see Table S1) in July and August 2019 on two reef zones: the back reef (1-2 m depth) and fore reef (5-10 m depth), between LTER sites 1 and 2 of the Mo’orea Coral Reef (MCR) Long Term Ecological Research (LTER) site. We selected fish species that broadly differ in their level of corallivory (Ezzat et al., 2020; Harmelin-Vivien and Bouchon-Navaro, 1983; Rotjan and Lewis, 2008; Viviani et al., 2019) as obligate corallivores (butterflyfishes *Chaetodon lunulatus*, *Chaetodon ornatissimus, Chaetodon reticulatus*, and the filefish *Amanses scopas*), facultative corallivores (butterflyfishes *Chaetodon citrinellus* and *Chaetodon pelewensis* and the parrotfish *Chlorurus spilurus*) and grazer/detritivores (surgeonfishes *Ctenochaetus flavicauda* and *Ctenochaetus striatus*). Additional details for the following sections are provided in the **Supplementary Methods**.

### Sample processing

Following collection, all samples were sub-sampled and processed for Symbiodiniaceae density, viability, and community composition analyses (except for corals, which were processed for Symbiodiniaceae community composition only). Characteristics of fecal material were also measured to support reef-scale estimates of Symbiodiniaceae cell dispersal. Feces were sampled from the hindgut of each fish (n=6-14 per species) so that Symbiodiniaceae condition (live versus dead) was assessed from cells that had passed through the entire fish digestive tract. Fecal samples for Symbiodiniaceae cell counts were preserved in 750 µl 10% formalin in 100 kDa-filtered seawater immediately after dissection out of each fish. Samples for DNA extractions were preserved in DNA/RNA shield (Zymo Research, CA). To calculate fecal densities for reef scale dispersal estimates, another fecal sub-sample was weighed and then its volume measured.

Samples of the common and abundant Mo’orean corals (Burkepile et al., 2019) *Acropora hyacinthus*, *Pocillopora* species complex (Gélin et al., 2017), and *Porites lobata* species complex (Forsman et al., 2015, 2009), were collected from fore reef and back reef sites (n=6 per reef zone). All coral samples were taken at least 5 meters apart. Each coral fragment was preserved for DNA extractions in DNA/RNA shield (Zymo Research, CA).

Reef-associated seawater samples were collected from the fore reef and back reef (n=6 per reef zone) at a distance of 10-100 cm off the reef bottom (at least 10 meters apart) and processed using a protocol modified from Littman et al (2008). Samples were filtered onto two separate 0.45-micron filters. One filter per sample was preserved for Symbiodiniaceae cell counts by resuspension in 3ml 5% formalin in 100 kDa-filtered seawater and vortexing at maximum speed for 10 minutes. The other filter was preserved for the isolation of genomic material in DNA/RNA shield.

Reef sediment samples were collected concomitantly with water column samples (n=6 per reef zone). Considering that Symbiodiniaceae populations in sediments are spatially heterogeneous (Littman et al., 2008), each individual sediment sample consisted of 500 ml of sediment pooled from ten sterile 50 ml conical tubes filled at five random locations in a 10m radius from where the associated seawater sample was collected. To best approximate the Symbiodiniaceae cells that would be available for uptake by nearby coral colonies, sediment collections were focused at the sediment-water interface (no deeper than 5 cm) and occurred less than 1m from live coral colonies. Sediments were immediately washed over a 120-micron nylon mesh using 500 ml 100 KDa filtered seawater and filtered through a 20-micron nylon mesh. The flow-through was then processed in the same manner as described above for the seawater samples.

### Symbiodiniaceae cell density and viability

All cell count samples were homogenized using a Fisher Scientific homogenizer F150 for five seconds to break up cell clumps and subsequently filtered through 120 and 20 micron nylon filters to reduce the density of debris in samples. Symbiodiniaceae cells in fish feces, seawater and sediments were quantified using a Neubauer hemocytometer. All samples were stained with 0.16% trypan blue 5 minutes before processing to parse live from dead cells (Haslun et al., 2011; Strychar and Coates, 2004; Zhang et al., 2008). Individual cells were interpreted as having been ‘live’ at the time of fixation if, following staining, they retained a golden-brown color (also see figure S1 and “Testing the efficacy of trypan blue for Symbiodiniaceae” in the Supplementary Materials). Cells were interpreted as ‘dead’ at the time of fixation if staining turned them blue. During counts, cells were counted as Symbiodiniaceae if they were of the correct size (6-12 µm; LaJeunesse et al., 2018), had visible organelles, a coccoid shape, and a large pyrenoid accumulation body (Littman et al., 2008; Nitschke et al., 2016).

We utilized cell culturing to confirm that Symbiodiniaceae cells in fish feces could remain viable for an extended period following passage through a consumer digestive tract culturing (n=61 fish fecal pellets, 24 replicate wells per sample; 1464 wells total; Table S1). Briefly, subsamples of fecal pellets were suspended in 1 ml of 100KDa-filtered seawater in a sterile petri dish. Then, 20 ul of this suspension was pipetted into each of 24 replicate wells in a 96-well plate filled with 200 ul sterile F/2 media. All culture wells were examined after 6-10 weeks using a compound microscope using with 20x and 40x magnification and scored for the presence/absence of intact Symbiodiniaceae-like cells.

### Symbiodiniaceae community composition

Genomic material was extracted using the ZymoBIOMICs DNA/RNA Miniprep kit (Zymo Research, CA), and the internal transcribed spacer-2 (ITS-2) region of Symbiodiniaceae rDNA was sequenced following Howe-Kerr et al (2020) using the primers SYM_VAR_5.8SII and SYM_VAR_REV (Hume et al., 2018). The resulting sequencing data were processed using Symportal (Hume et al., 2019). Samples with <1,000 reads were discarded and sequencing depth was assessed using rarefaction curves. To circumvent issues related to the interpretation of inter-versus intragenomic variation in the Symbiodiniaceae ITS-2 region (Correa et al., 2009; LaJeunesse and Pinzón, 2007), ITS2-profiles were reduced to number of reads per Symbiodiniaceae genus and expressed as percentages.

### Reef-scale Symbiodiniaceae dispersal estimates

We estimated the number of Symbiodiniaceae cells dispersed by three species of corallivorous fish per 100 m^2^ per day by applying a bootstrap approach (1000 iterations) to the equation *T* = *eSWCF*, where the estimated number of dispersed Symbiodiniaceae cells (*T*) is the product of five variables: A species-specific constant representing the estimated number of egestions in an eight-hour day (*e*); fecal pellet sizes in cm (*S*); densities of fecal samples in g cm^−1^ (*W*); densities of Symbiodiniaceae cells g^−1^ feces (*C*); and fish densities 100 m^−2^ (*F*; extracted from the MCR LTER dataset http://mcrlter.msi.ucsb.edu/cgi-bin/showDataset.cgi?docid=knb-lter-mcr.6; accessed February 14, 2020).

We collected data on fish egestion rates (*e*) and fecal pellet sizes (*S*) *in situ* via fish follows in the field between 09:00h and 17:00h (Table S6). Individuals of C*. ornatissimus* (n=53), *C. reticulatus* (n=39) and *C. citrinellus* (n=31), were each followed for a mean of 7.9 minutes (16.2 hours of observations total) and all egestion events were recorded. Estimated egestion rates per hour were calculated as the total number of observed egestions divided by the total period of observations per species. We conservatively assumed that the observed fish species were active for eight hours per day, equal to the time frame over which we made our observations, and that egestion rates remained constant throughout the day. We measured the length of any fecal pellet that fell on a flat surface and remained intact (N=12) using a standard ruler. To convert the lengths of fecal pellets to weights (*W*), we used a precision balance to weigh pre-measured fecal samples from collected individuals of *C. citrinellus* (n=4), *C. ornatissimus* (n=3) and *C. reticulatus* (n=3). Cell densities (*C*) calculated as described above were expressed as live cells g^−1^ feces.

### Statistical Analyses

For all tests, biological replicates were defined as samples collected from distinct fish individuals; or samples from coral colonies separated by at least 5m; or sediment or water samples collected at least 10m apart. Technical replicates (e.g., in Symbiodiniaceae cell counts) were defined as analyses conducted on multiple sub-samples of an individual biological replicate and not included in the statistical analysis.

We tested for differences in Symbiodiniaceae cell densities among sample categories (obligate corallivores, facultative corallivores, grazer/detritivores and sediment and seawater) using Kruskal-Wallis tests and Dunn tests with Bonferroni-corrected p-values for multiple comparisons (Tables S2&S3) using dunn.test package v1.3.5. in R version 3.6.1 (R Core Team, 2019). Tests among individual fish species and sediment and seawater samples were conducted in the same way. We used non-parametric tests because linear models did not follow the model assumptions (even after transformation), but non-parametric and parametric tests gave the same general results.

We used a PERMANOVA in Vegan 2.5-6 (Oksanen et al., 2019) to test for differences in genus-level Symbiodiniaceae community compositions among obligate corallivores, facultative corallivores, grazer/detritivores, individual coral species, and sediments and seawater. However, we found significant heterogeneous dispersion between sample categories using betadisper(). Because PERMANOVA is sensitive to heterogeneous dispersion in concert with an unbalanced design (Anderson and Walsh, 2013), we randomly subsampled 12 rows for each of the four sample categories (see **Supplementary Methods** and **Figure 3–Source Data 1** for sample names). We then tested for differences between sample categories using PERMANOVA with adonis() based on untransformed Bray-Curtis distances. Pairwise tests were conducted with a Benjamini-Hochberg error correction using pairwiseAdonis 0.4 (Martinez Arbizu, 2020).

## Acknowledgements

Special thanks go to Deron Burkepile and Erika Eliason for input on the study design and providing some materials related to this work. For support in the field, the authors express their sincere appreciation to Jake Emmert and Ryan Hannum from the Aquarium at Moody Gardens in Galveston, Texas, and to Journ Galvan, Haley Glasman, Maya Gorgas, Ashtyn Isaak, Kai Kopecky, Kaitlyn Landfield, Hunter Lenihan, Jacey van Wert and Erin Winslow. Further thanks go out to Tom E.X. Miller, Dave W. Armitage and Daniel Gorczynski for input on the data analysis. Lastly, many thanks to Janavi Mahimtura Folmsbee for designing and creating a painting for figure 1, and Thiago B.S. Correa for a suggestion that improved figure 1. This work represents a contribution of the Moorea Coral Reef (MCR) LTER Site (NSF OCE 16–37396).

## Figure Supplements

**Figure 2 – figure supplement 1.**
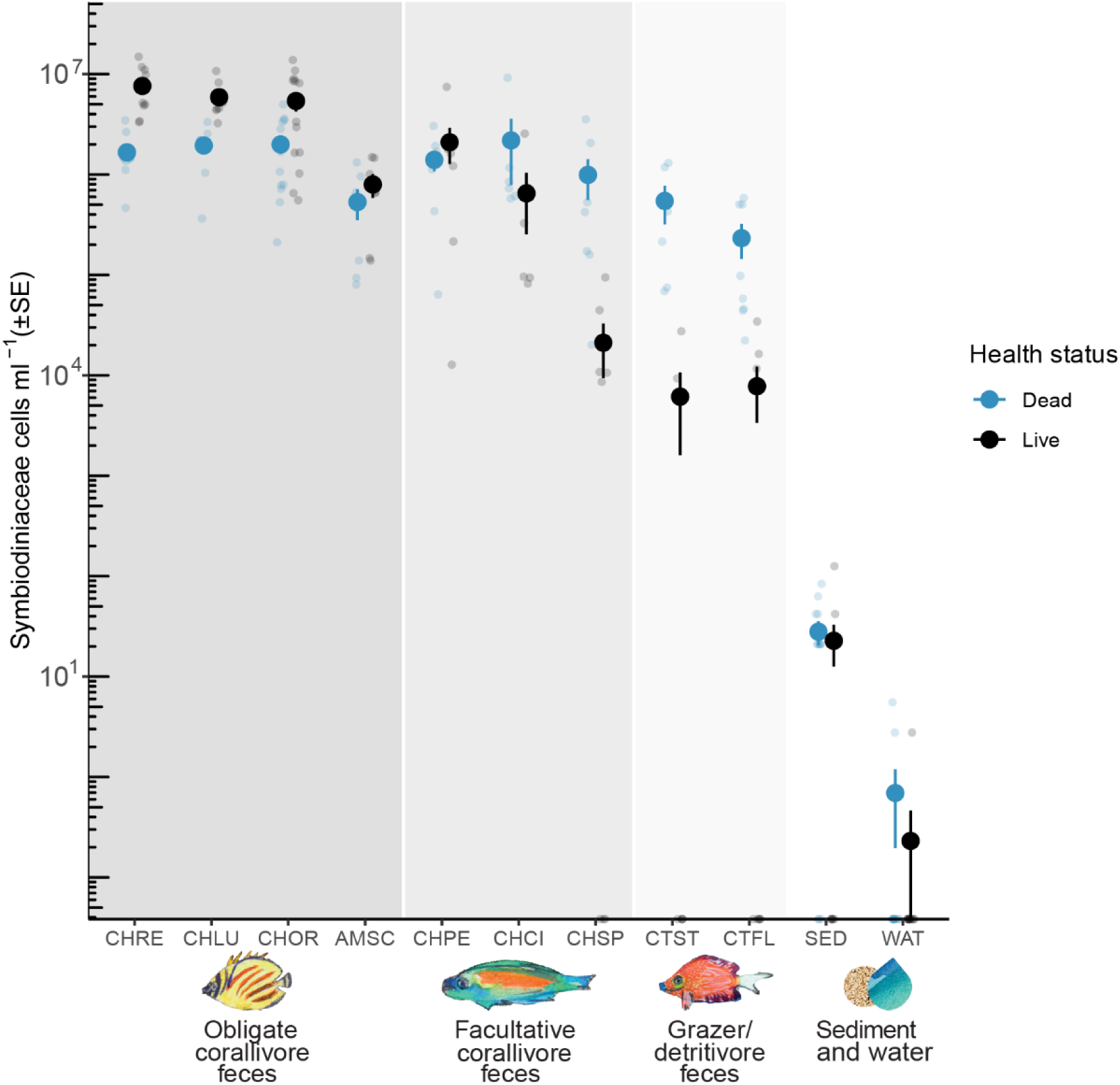
Densities of live (black) and dead (blue) Symbiodiniaceae cells per sample type. Large dots indicate the mean of a given sample type; vertical lines depict the standard error. Small blue and black dots indicate values from individual samples for dead and live Symbiodiniaceae cell densities, respectively. Included samples (and sample sizes) for each sample category are as follows: Obligate corallivores (dark shading): Amanses scopas, AMSC (7); Chaetodon lunulatus, CHLU (8); Chaetodon ornatissimus, CHOR (14); Chaetodon reticulatus, CHRE (11). Facultative corallivores (intermediate shading): Chaetodon pelewensis, CHPE (8); Chaetodon citrinellus, CHCI (6); and Chlorurus spilurus CHSP (8). Grazer/Detritivores (light shading): Ctenochaetus flavicauda, CTFL (8); and Ctenochaetus striatus, CTST (6). Sediment and water (no shading): Sediment, SED (12); Water, WAT (12). Significant differences are listed in Table S3. For data used see **Figure 2-Source Data 1**.

## Source Data

### Figure 2 – Source Data 1

The file “Fishces_2019_cellcounts.csv” contains densities of live (unstained by trypan blue; column ‘Live’) and dead (stained by trypan blue; column ‘Dead’) Symbiodiniaceae cells ml^−1^ material (feces, sediment seawater), as determined through cell counts using a hemocytometer and compound microscope at 20x and 40x magnification. Reported cell densities are averages of 8 replicate counts. The code to replicate this analysis can be found on https://github.com/CorreaLab/Fishces_2020/blob/master/Symbiodiniaceae%20cell%20counts%20code.

### Figure 3 – Source Data 1

Sequencing data after processing in Symportal (Hume et al., 2019). Individual ITS-2 profiles have been removed to facilitate analysis of Symbiodiniaceae communities at the genus level. All. csv files in the folder “Figure3-source_data_1.zip” are required for analysis using code on https://github.com/CorreaLab/Fishces_2020/blob/master/Symbiodiniaceae%20community%20analysis%20code. The file “Random_subsampling_data.xlsx” contains the randomly subsampled data used for statistical analysis with PERMANOVA.

### Figure 4 – Source Data 1

The file “Bootstrap_datafile_manuscript.csv” contains the dataset used to estimate the daily dispersal of Symbiodiniaceae cells by three fish species. the column ‘live_g_sample’ contains Symbiodiniaceae cell densities g^−1^ fecal sample; the column ‘wet weight’ contains the weight of fecal pellets cm^−1^; the columns ‘Site’ – ‘Density’ contain fish abundances at four transects of MCR LTER sites 1 and 2; the column ‘Pellet_length’ contains lengths of fecal pellets as measured during observations in the field. The code to replicate this analysis can be found on https://github.com/CorreaLab/Fishces_2020/blob/master/Symbiodiniaceae%20cell%20dispersal%20bootstrap%20code.

## Supplementary Materials

### Supplementary Methods

Individuals of nine fish species (n=6-14 per sample type or species, see Table S1), as well as corals, sediments and seawater were collected in July and August 2019 on the back reef (1-2 m depth) and fore reef (5-10 m depth), between LTER sites 1 and 2 of the Mo’orea Coral Reef (MCR) Long Term Ecological Research (LTER) site. We selected fish species that broadly differ in their level of corallivory (Ezzat et al., 2020; Harmelin-Vivien and Bouchon-Navaro, 1983; Rotjan and Lewis, 2008; Viviani et al., 2019) as obligate corallivores (butterflyfishes *Chaetodon lunulatus*, *Chaetodon ornatissimus, Chaetodon reticulatus*, and the filefish *Amanses scopas*), facultative corallivores (butterflyfishes *Chaetodon citrinellus* and *Chaetodon pelewensis* and the parrotfish *Chlorurus spilurus*) and grazer/detritivores (surgeonfishes *Ctenochaetus flavicauda* and *Ctenochaetus striatus*).

#### Sample processing overview

Following collection, all samples were sub-sampled and processed for Symbiodiniaceae density, viability and community composition analyses (except for corals, which were processed for Symbiodiniaceae genetic diversity only). Characteristics of fecal material (density, weight) were also measured to support reef-scale estimates of Symbiodiniaceae cell dispersal.

##### Fish fecal samples

Feces were sampled from the hindgut of each fish so that Symbiodiniaceae condition (live versus dead) was assessed from cells that had passed through the entire fish digestive tract. Fecal samples for Symbiodiniaceae cell counts were preserved in 750 µl 10% formalin in 100 kDa-filtered seawater immediately after dissection. Collection tubes were weighed before and after sample addition using a 1 mg precision balance to determine the sample weight. Sample weights ranged from 22 to 170 mg (μ=73, σ= 32). Sample volumes were measured after resuspension in the fixative using a 1000p pipette and subtracting the volume of fixative. Sample volumes ranged from 42 to 234 μl (μ=109, σ= 45). A similarly sized sample of feces was placed in DNA/RNA shield Lysis and Collection tubes (Zymo Research, CA) for extraction of genomic material. Lastly, if enough sample was available, another subsample was transferred to a sterile 1.5 ml tube, weighed, and processed for culturing of Symbiodiniaceae cells.

##### Coral samples

Coral samples were collected from fore reef and back reef sites (n=6 per species at each sites). We sampled coral species that are common in Mo’orea: *Acropora hyacinthus*, *Pocillopora* species complex (Gélin et al., 2017), and *Porites lobata* species complex (Forsman et al., 2015, 2009). All coral samples were taken at least 5 meters apart. Each coral fragment was placed in 6 ml of DNA/RNA shield (Zymo Research, CA) with 1.35g Lysing Matrix A garnet and 5 0.6-cm ceramic beads (MPBio, California) and vortexed for 20 minutes for extraction of genomic material.

##### Seawater samples

Seawater samples were collected from the fore reef and back reef (n=6 per reef zone and sample type) and processed using a protocol modified from Littman et al. (2008). Briefly, 3.8 L water samples were collected at least 10 meters apart in a polyethylene jug 10-100cm above the substrate and transported to the molecular lab at the Richard B. Gump research station. The samples were concentrated to ~100 ml using tangential flow filtration (0.2 micron), divided into two 50 ml subsamples and filtered onto two separate 0.45-micron filters using a vacuum pump. One filter per sample was preserved for Symbiodiniaceae cell counts by resuspension in 3ml 5% formalin in 100 kDa-filtered seawater and vortexing at maximum speed for 10 minutes. The other filter was preserved for the isolation of genomic material by placing it in 5 ml of DNA/RNA shield (Zymo Research, CA) with 1.35g Lysing Matrix A garnet and 5 0.6-cm ceramic beads (MPBio, California) and vortexed for 20 minutes for extraction of genomic material.

##### Sediment samples

Reef sediment samples were collected concomitantly with water column samples. Considering that Symbiodiniaceae populations in sediments are spatially heterogeneous (Littman et al., 2008), each individual sediment sample consisted of 500 ml of sediment pooled from ten sterile 50 ml conical tubes filled at five random locations in a 10m radius from where the associated water sample was collected. All sediment samples were taken at least 10 meters apart. To best approximate the Symbiodiniaceae cells that would be available for uptake by nearby coral colonies, sediment collections were focused at the sediment-water interface (no deeper than 5 cm). All sediment samples were collected less than a meter from live coral colonies. At the lab, sediments were immediately washed over a 120-micron nylon mesh using 500 ml 100 KDa filtered seawater and filtered through a 20-micron nylon mesh. The flow-through was then processed in the same manner as described above for the seawater samples.

#### Testing the efficacy of trypan blue for Symbiodiniaceae

Trypan blue staining is an established method for parsing live from dead cells in formaldehyde-fixed mammalian cells (Zhang et al., 2008) and in non-fixed Symbiodiniaceae cells (e.g. Haslun et al., 2011; Strychar and Coates, 2004), but has to our knowledge not been applied to formalin-fixed Symbiodiniaceae cells. To test whether fixing samples of Symbiodiniaceae cells 10% formalin interferes with the efficacy of trypan blue in staining dead Symbiodiniaceae cells, we conducted a replicated experiment using Symbiodiniaceae cultures (Figure S1). Briefly, three different Symbiodiniaceae strains (*Breviolum* sp. *Mf1.5b, Symbiodinium microadriaticum*, and *Fugacium kawagutii*) were sampled (1 ml each) and half of each sample was kept under control conditions or heat-killed at 80°C for one hour, similar to (Franklin et al., 2006). To test the effect of sample fixation on the efficacy of trypan blue - as well as the order in which samples were fixed and stained - samples were then either: immediately fixed in 10% formalin and stained with trypan blue after 48 hours (FOTB+); first stained with trypan blue and fixed with 10% formalin after five minutes (TBFO); fixed with 10% formalin and stained with trypan blue immediately after (FOTB); or stained with trypan blue and not fixed (TB). Fixation, staining and hemocytometry proceeded as detailed in the next section “Symbiodiniaceae cell density and viability”. We expected that heat-killed Symbiodiniaceae cells with compromised membrane integrity would be stained blue in all fixation treatments, while intact cells would retain their unstained golden-brown color (Haslun et al., 2011; Strychar and Coates, 2004). In our experiment, the fraction of live, non-stained Symbiodiniaceae cells differed between the heat and control treatment (two-way ANOVA: df=1, F=1263.473, p<0.001) but not between staining and preservation methods (df=3, F=0.209, p=0.889, Figure S1). These results indicate that preservation with 10% formalin does not inhibit the efficacy of trypan blue in staining Symbiodiniaceae cells that were dead at the time of fixation.

#### Symbiodiniaceae cell density and viability

All samples were homogenized using a Fisher Scientific homogenizer F150 for five seconds to break up cell clumps. Samples were then filtered through 120 and 20 micron nylon filters to reduce the density of debris in samples. Next, samples were stained with 0.4% trypan blue at a 2:5 ratio for a final concentration of 0.16% 5 minutes before processing. After trypan blue staining and preservation, hemocytometry was conducted using a light microscope with 20X magnification and the cell density per 0.1 µl fixative was determined by averaging counts from 8 separate 1mm^2^ squares (8 technical replicates). Cells were counted as Symbiodiniaceae if they were of the correct size (6-12 µm; LaJeunesse et al., 2018), had visible organelles, a coccoid shape, and a large pyrenoid accumulation body (Littman et al., 2008; Nitschke et al., 2016). Subsamples were frequently viewed at higher magnification to verify their identity, using cultured Symbiodiniaceae cells (live and dead) as visual references. We expressed cell densities as cells per ml source material (feces, water, sediment) by dividing the concentration by the fraction of sample in the summed volume of fixative, stain and sample volume, and multiplying this by 1,000.

#### Symbiodiniaceae culturing

When sufficient fecal material was available (N=61; Table S1), we utilized cell culturing to confirm that Symbiodiniaceae cells can remain viable for an extended period following passage through a corallivore digestive tract. Briefly, subsamples of fecal pellets between 8 and 153 mg (μ=40, σ= 27) were suspended in 1 ml of 100KDa-filtered seawater in a sterile petri dish. Then, 20 µl of this suspension was pipetted into each of 24 replicate wells in a 96-well plate filled with 200 ul sterile F/2 media. The culture plates were stored at room temperature under fluorescent lighting on a ~12:12 light:dark cycle for up to six weeks and then transported back to Rice University (Houston, TX, U.S.A.), where they were transferred to an incubator set to 25°C on a 12:12 light:dark cycle. All wells were examined within ten weeks of initiating the cultures using a compound microscope with 20x and 40x magnification; each well was scored for the presence/absence of intact Symbiodiniaceae cells, based on the cell characteristics listed previously in this section.

#### Symbiodiniaceae community composition

DNA was extracted using the ZymoBIOMICs DNA/RNA Miniprep kit shield (Zymo Research, CA). The ITS-2 region of Symbiodiniaceae rDNA was amplified using SYM_VAR_5.8SII (5’ GAATTGCAGAACTCCGTGAACC 3’) and SYM_VAR_REV (5’ CGGGTTCWCTTGTYTGACTTCATGC 3’) primers (Hume et al., 2018) and sequenced on the Illumina MiSeq platform at the Georgia Genomics and Bioinformatics Core (University of Georgia, Athens, GA) following details outlined in Howe-Kerr et al. (2020). Demultiplexed fastq files were generated with Illumina’s BaseSpaceFS (version 1.5.964) and fastq files were processed using Symportal (Hume et al., 2019). Symportal is an analytical framework that resolves Symbiodiniaceae taxa using NGS data of the ITS-2 marker gene by separating reads into different Symbiodiniaceae genera and collapsing the intragenomic variation of the ITS-2 gene by assigning ITS-2 profiles (Hume et al., 2019). Samples with <1,000 reads were discarded and sequencing depth was assessed using rarefaction curves. The resulting ITS2-profiles were reduced to number of reads per Symbiodiniaceae genus and expressed as percentages.

#### Reef-scale Symbiodiniaceae dispersal estimates

We estimated the number of Symbiodiniaceae cells dispersed by the obligate corallivores *C. ornatissimus* and *C. reticulatus* and the facultative corallivore *C. citrinellus* per 100 m^2^ per day by applying a bootstrap approach (1000 iterations) to the following equation:

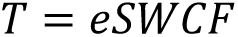

Here, the estimated number of dispersed Symbiodiniaceae cells (*T*) is the product of five variables: A fish species-specific constant representing the estimated number of egestions (*e*) in an eight-hour day based on species-specific egestion rates; fecal pellet sizes in cm (*S*); densities of fecal samples in g cm^−1^ (*W*); densities of Symbiodiniaceae cells g^−1^ feces (*C*); and fish densities per 100 m^2^ (*F*).

Dispersal rates were calculated for the fore reef for butterflyfishes *Chaetodon ornatissimus* and *Chaetodon reticulatus* and for the back reef for *Chaetodon citrinellus* (Table S6). For *C. ornatissimus* and *C. reticulatus*, three of the variables (*e, S*, and *W*) utilized data pooled from the fore reef and back reef, whereas data for the two other variables (*C* and *F*) are from the fore reef only. For *C. citrinellus*, data for all five variables are from the back reef.

We collected data on fish egestion rates (*e*) and fecal pellet sizes (*S*) *in situ* via fish follows in the field. All follows were conducted between 09:00 and 17:00 at the same sites where we collected fish, seawater and sediment samples. Individuals of C*. ornatissimus* (n=53), *C. reticulatus* (n=39) and *C. citrinellus* (n=31), were each followed for a mean of 7.9 minutes (16.2 hours of observations total) and all egestion events were recorded. If the same individual fish egested twice within a single follow, this was interpreted and recorded as part of a single egestion event. Estimated egestion rates per hour were calculated as the total number of observed egestions divided by the total period of observations per species (Table S6): *C. ornatissimus*, n=53, 7.4 hours total observations, 10 observed egestions; *C. reticulatus*, n=39, 4.6 hours total observations, 5 observed egestions; *C. citrinellus*, n=31, 4.1 hours total observations, 4 observed egestions. We conservatively assumed that the observed fish species were active for eight hours per day, equal to the time frame over which we made our observations and that egestion rates remained constant throughout the day. We measured the length of any fecal pellet that fell on a flat surface and remained intact (N=12) using a standard ruler (Table S6). To convert the lengths of fecal pellets to weights (*W*), we measured the length of individual fecal samples collected from *C. citrinellus* (n=4), *C. ornatissimus* (n=3) and *C. reticulatus* (n=3) and then used a precision balance to weigh each of these feces. Cell densities (*C*) calculated as described above were expressed as cells g^−1^ feces.

Fish densities (*F*) were extracted from the MCR LTER dataset (http://mcrlter.msi.ucsb.edu/cgi-bin/showDataset.cgi?docid=knb-lter-mcr.6 accessed February 14, 2020). We used fish densities from the fore reef for *Chaetodon ornatissimus* and *Chaetodon reticulatus* and data from back reef sites for *Chaetodon citrinellus*, because the latter species does not occur on the fore reef. Data from 2019 were not available as of April 2020, so we therefore used data from 2018. We did not use data from previous years because corallivorous fish populations have been recovering from a population crash since 2010, that was caused by a crown of thorns starfish outbreak (2006-2010) and Cyclone Oli (2010; Adjeroud et al., 2018). Briefly, the MCR LTER collects these data by counting fishes in four replicate 5×50 m fish belt transects at the two north-shore LTER sites (LTER 1 and 2), resulting in densities of fish 250m^−2^. In this study, we multiplied the LTER data by 0.4 to convert densities to 100m^−2^.

#### Coral cover transects

Replicate line transects were conducted at our fore reef (n=5) and back reef sites (n=12) to determine live coral cover. Briefly, a 50m transect was rolled out parallel to the shoreline and pictures were taken to categorize substrates as “live coral” or “other” at each meter mark.

#### Statistical analyses

For all tests, biological replicates were defined as samples collected from distinct fish individuals; or samples from coral colonies separated by at least 5m; or sediment or water samples collected at least 10m apart (see supplemental methods). Technical replicates (e.g., in Symbiodiniaceae cell counts) were defined as analyses conducted on multiple sub-samples of an individual biological replicate and not included in the statistical analysis.

We tested for differences in Symbiodiniaceae cell densities among sample groups (obligate corallivores, facultative corallivores, grazer/detritivores and sediment and seawater) using Kruskal-Wallis tests and Dunn tests with Bonferroni-corrected p-values for multiple comparisons (Tables S2&S3) using dunn.test package v1.3.5. in R version 3.6.1 (R Core Team, 2019). Tests among individual fish species, and sediment and seawater samples were conducted in the same way. Individual samples (feces, sediment or water) were considered as biological replicates. We checked for outliers using boxplots and observed one outlier among the facultative corallivore samples. However, this datapoint was in the same range as data for obligate corallivores and was therefore retained. We used non-parametric tests because linear models did not follow the model assumptions (even after transformations); non-parametric and parametric tests gave the same results.

We used a PERMANOVA in Vegan 2.5-6 (Oksanen et al., 2019) to test for differences in genus-level Symbiodiniaceae community compositions among sample categories (obligate corallivores, facultative corallivores, grazer/detritivores, individual coral species, and sediments and seawater). Individual samples (feces, sediment or water) were considered as biological replicates for each sample category. We found significant heterogeneous dispersion between sample categories using betadisper(). Because PERMANOVA is sensitive to heterogeneous dispersion in concert with an unbalanced design (Anderson and Walsh, 2013), we used sample_n() to randomly subsampled 12 rows for each of the four sample categories (See **Figure 3-Source Data 1**: All coral samples; Obligate corallivores: 19AMSC4, 19AMSC6, 19AMSC7, 19CHLU2, 19CHLU6, 19CHOR5, 19CHOR7, 19CHOR8, 19CHOR12, 19CHOR14, 19CHRE9, 19CHRE11; facultative corallivores: 19CHCI2, 19CHCI3, 19CHCI4, 19CHPE1, 19CHPE3, 19CHPE4, 19CHPE5, 19CHPE6, 19CHPE8, 19CHSP2, 19CHSP4, 19CHSP7; Grazer/detritivores: 19CTFL1, 19CTFL3, 19CTFL4, 19CTFL5, 19CTFL6, 19CTFL7, 19CTFL8, 19CTST1, 19CTST3, 19CTST4, 19CTST5, 19CTST6; sediment and water: BAKSED1, BAKSED2, BAKSED4, BAKSED5, FORSED1, FORSED2, FORSED3, FORSED4, FORSED6, FORH2O2, BAKH2O5, FORH2O5). We then tested for differences between sample categories using PERMANOVA with adonis() using Bray-Curtis distances. Pairwise tests were conducted with a Benjamini-Hochberg error correction using pairwiseAdonis 0.4 (Martinez Arbizu, 2020).

To test for differences in live coral cover between the back reef and fore reef zones, we used an analysis of variance (ANOVA). We tested for normality of the distributions and of the residuals using a Shapiro test. The assumption of homogeneity of variance was tested using Levene’s test.

---

**Table S1:** Summary of the samples collected and processed in this study for Symbiodiniaceae cell density and viability (Figure 2), culturing, and community composition based on the internal transcribed spacer-2 (ITS-2) region of rDNA (Figure 3). *Pocillopora* spp. = *Pocillopora* species complex; *Porites lobata* spp. = *Porites lobata* species complex.

| Sample category and type | Code | Symbiodiniaceae cell density & viability |  |  | Symbiodiniaceae culturing |  |  | Symbiodiniaceae community composition (>1,000 ITS-2 reads) |  |  |
| --- | --- | --- | --- | --- | --- | --- | --- | --- | --- | --- |
|  |  | Back reef | Fore reef | total | Back reef | Fore reef | total | Back reef | Fore reef | total |
| Obligate corallivores |  |  |  |  |  |  |  |  |  |  |
| *Amanes scopas* | AMSC | 0 | 7 | 7 | 0 | 6 | 6 | 0 | 7 | 7 |
| *Chaetodon lunulatus* | CHLU | 7 | 1 | 8 | 5 | 1 | 6 | 7 | 1 | 8 |
| *Chaetodon ornatissimus* | CHOR | 6 | 8 | 14 | 6 | 7 | 13 | 6 | 8 | 14 |
| *Chaetodon reticulatus* | CHRE | 3 | 8 | 11 | 3 | 7 | 10 | 3 | 8 | 11 |
| <u>Category sum</u> |  | <u>16</u> | <u>24</u> | <u>40</u> | <u>14</u> | <u>21</u> | <u>35</u> | <u>16</u> | <u>24</u> | <u>40</u> |
| Facultative corallivores |  |  |  |  |  |  |  |  |  |  |
| *Chaetodon citrinellus* | CHCI | 6 | 0 | 6 | 6 | 0 | 6 | 6 | 0 | 6 |
| *Chaetodon pelewensis* | CHPE | 0 | 8 | 8 | 0 | 4 | 4 | 0 | 8 | 8 |
| *Chlorurus spilurus* | CHSP | 0 | 8 | 8 | 0 | 8 | 8 | 0 | 8 | 8 |
| <u>Category sum</u> |  | <u>6</u> | <u>16</u> | <u>22</u> | <u>6</u> | <u>12</u> | <u>18</u> | <u>6</u> | <u>16</u> | <u>22</u> |
| Grazer/detritivores |  |  |  |  |  |  |  |  |  |  |
| *Ctenochaetus flavicauda* | CTFL | 0 | 8 | 8 | 0 | 8 | 8 | 0 | 7 | 7 |
| *Ctenochaetus striatus* | CTST | 6 | 0 | 6 | - | - | - | 6 | 0 | 6 |
| <u>Category sum</u> |  | <u>6</u> | <u>8</u> | <u>14</u> | <u>0</u> | <u>8</u> | <u>8</u> | <u>6</u> | <u>7</u> | <u>13</u> |
| Sediment and water |  |  |  |  |  |  |  |  |  |  |
| Sediment | SED | 6 | 6 | 12 | - | - | - | 6 | 6 | 12 |
| Water | WAT | 6 | 6 | 12 | - | - | - | 3 | 4 | 7 |
| <u>Category sum</u> |  | <u>12</u> | <u>12</u> | <u>24</u> | - | - | - | <u>9</u> | <u>10</u> | <u>19</u> |
| Corals |  |  |  |  |  |  |  |  |  |  |
| *Acropora hyacinthus* | ACR | - | - | - | - | - | - | 6 | 5 | 11 |
| *Pocillopora* spp. | POC | - | - | - | - | - | - | 6 | 6 | 12 |
| *Porites lobata* spp. | POR | - | - | - | - | - | - | 6 | 6 | 12 |
| <u>Category sum</u> |  | - | - | - | - | - | - | <u>18</u> | <u>17</u> | <u>35</u> |

**Figure S1:**
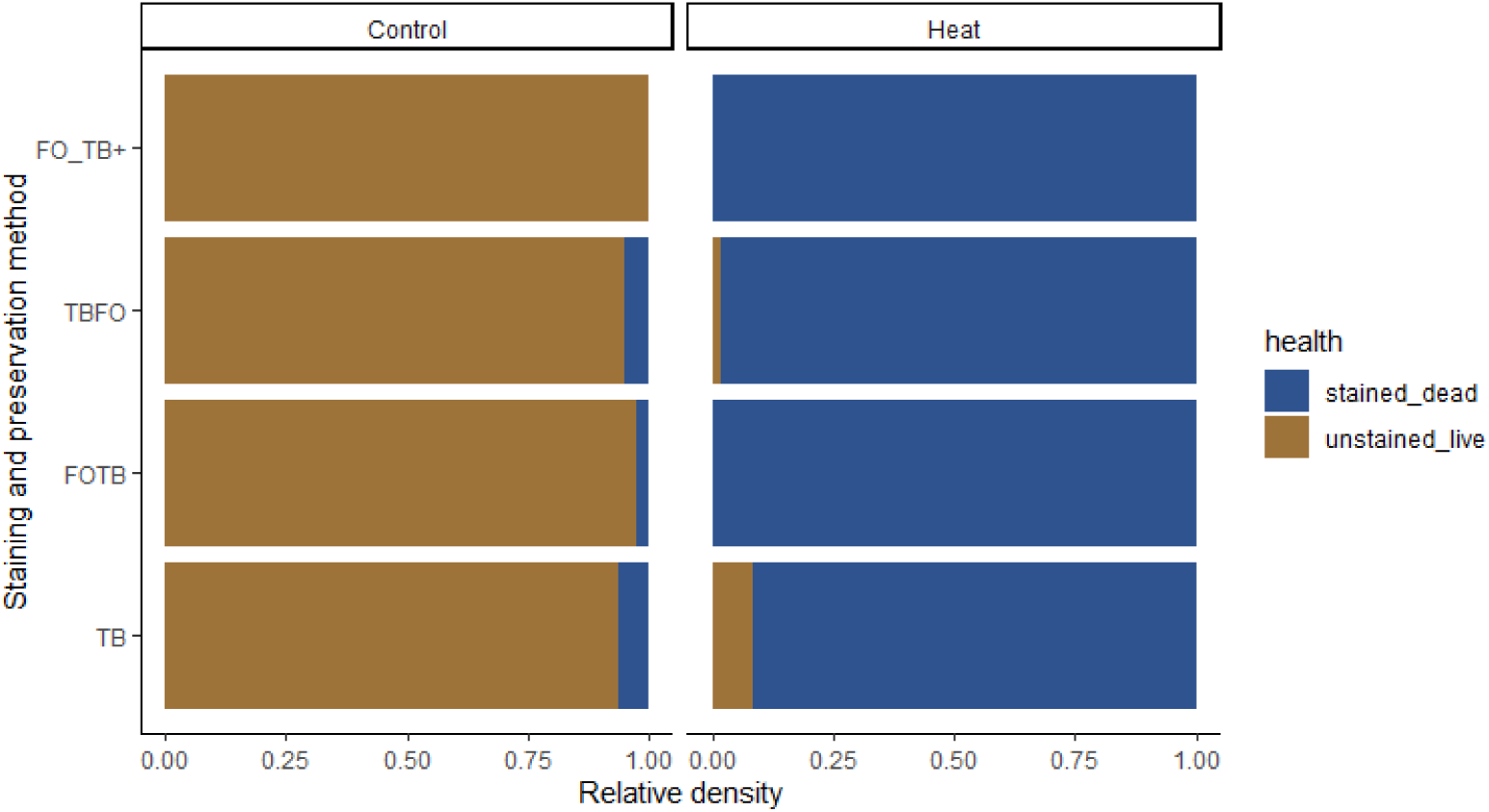
Hemocytometry with trypan blue stain accurately differentiates live and dead Symbiodiniaceae cells that were fixed in 10% formalin. We conducted a replicated experiment (see Supplementary Methods section ‘Testing the efficacy of trypan blue for Symbiodiniaceae’) to confirm that hemocytometry of formalin-fixed samples in conjunction with the trypan blue stain assay accurately parses dead and live Symbiodiniaceae cells. Cultures of three Symbiodiniaceae strains (*Breviolum* sp. *Mf1.5b, Symbiodinium microadriaticum*, and *Fugacium kawagutii*) were sampled and half of each sample was kept at room temperature (Control) or exposed to 80°C for one hour (Heat) similar to (Franklin et al., 2006). The following staining and preservation methods were tested: FO_TB+: immediately fixed in 10% formalin and stained with trypan Blue after 48 hours; TBFO: First stained with trypan blue and fixed with formalin after five minutes; FOTB: Fixed with 10% formalin and stained with trypan blue immediately afterward; TB: Stained with trypan blue and not fixed. The fraction of live Symbiodiniaceae cells differed between the heat and control treatment (two-way ANOVA: df=1, F=1263.5, p<0.001) but not between staining and preservation methods (df=3, F=0.2, p=0.889). These results indicate that preservation with 10% formalin and staining with trypan blue is an appropriate method for quantifying live and dead Symbiodiniaceae cells.

**Table S2:**
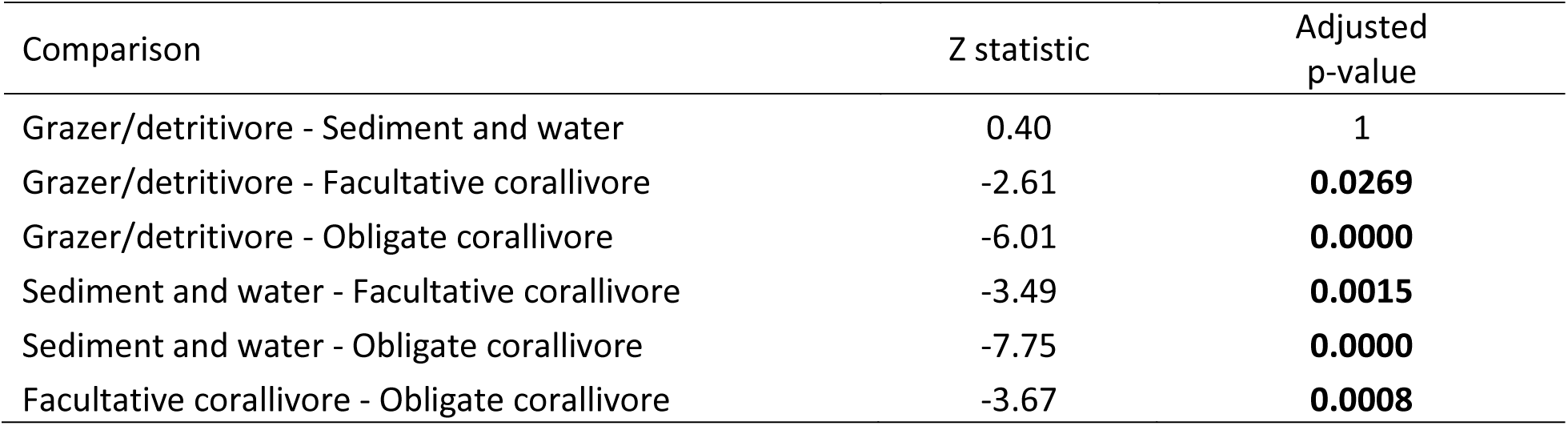
Pairwise differences in dinoflagellate symbiont cell densities between sample categories (Figure 2). Displayed are Dunn test results with Bonferroni-adjusted p-values. Significant p-values (<0.05) are bolded. Overall Kruskal-Wallis test results: Chi-squared=74.3248, df=3, p-value<0.001. Included samples (and sample sizes) for each sample category are as follows: Obligate corallivores: *Amanses scopas*, AMSC (7)*; Chaetodon lunulatus*, CHLU (8); *Chaetodon ornatissimus*, CHOR (14); *Chaetodon reticulatus*, CHRE (11). Facultative corallivores: *Chaetodon pelewensis*, CHPE (8); *Chaetodon citrinellus*, CHCI (6); and *Chlorurus spilurus* CHSP (8). Grazer/Detritivores: *Ctenochaetus flavicauda*, CTFL (8); and *Ctenochaetus striatus*, CTST (6). Sediment and water: Sediment, SED (12); Water, WAT (12).

**Table S3:** Results of pairwise Dunn tests on Symbiodiniaceae cell densities between species, sediment and water samples (Figure 2-figure supplement 1). We controlled for false positives with the Benjamini-Hochberg procedure. Significant p-values (<0.05) are bolded. Overall Kruskal-Wallis test results: chi-squared=85.2132, df=10, p-value=0. Included samples (and sample sizes) for each sample type are as follows: Obligate corallivores (Obl): *Amanses scopas*, AMSC (7)*; Chaetodon lunulatus*, CHLU (8); *Chaetodon ornatissimus*, CHOR (14); *Chaetodon reticulatus*, CHRE (11). Facultative corallivores (Fac): *Chaetodon pelewensis*, CHPE (8); *Chaetodon citrinellus*, CHCI (6); and *Chlorurus spilurus* CHSP (8). Grazer/Detritivores (Gra): *Ctenochaetus flavicauda*, CTFL (8); and *Ctenochaetus striatus*, CTST (6). Sediment and water (Env): Sediment, SED (12); Water, WAT (12).

| Species comparison | Species diets | Z statistic | Adjusted p-value |
| --- | --- | --- | --- |
| AMSC - CHLU | Obl - Obl | -1.67 | 0.0771 |
| AMSC - CHOR | Obl - Obl | -1.52 | 0.0978 |
| AMSC - CHRE | Obl - Obl | -2.10 | **0.0363** |
| CHLU - CHOR | Obl - Obl | 0.36 | 0.3961 |
| CHLU - CHRE | Obl - Obl | -0.33 | 0.4005 |
| CHOR - CHRE | Obl - Obl | -0.77 | 0.2880 |
| AMSC - CHCI | Obl - Fac | 0.19 | 0.4494 |
| AMSC - CHPE | Obl - Fac | -0.46 | 0.3640 |
| AMSC - CHSP | Obl - Fac | 1.75 | 0.0683 |
| CHLU - CHPE | Obl - Fac | 1.25 | 0.1516 |
| CHLU - CHSP | Obl - Fac | 3.54 | **0.0007** |
| CHOR - CHPE | Obl - Fac | 1.06 | 0.1996 |
| CHOR - CHSP | Obl - Fac | 3.64 | **0.0005** |
| CHRE - CHSP | Obl - Fac | 4.14 | **0.0001** |
| AMSC - WAT | Obl - Env | 3.09 | **0.0032** |
| AMSC - SED | Obl - Env | 2.39 | **0.0213** |
| CHLU - WAT | Obl - Env | 5.11 | **0.0000** |
| CHLU - SED | Obl - Env | 4.38 | **0.0000** |
| CHOR - WAT | Obl - Env | 5.53 | **0.0000** |
| CHOR - SED | Obl - Env | 4.68 | **0.0000** |
| CHRE - WAT | Obl - Env | 5.95 | **0.0000** |
| CHRE - SED | Obl - Env | 5.15 | **0.0000** |
| AMSC - CTFL | Obl - Gra | 2.20 | **0.0306** |
| AMSC - CTST | Obl - Gra | 2.16 | **0.0323** |
| CHLU - CTFL | Obl - Gra | 4.00 | **0.0002** |
| CHLU - CTST | Obl - Gra | 3.83 | **0.0003** |
| CHOR - CTFL | Obl - Gra | 4.16 | **0.0001** |
| CHOR - CTST | Obl - Gra | 3.91 | **0.0002** |
| CHRE - CTFL | Obl - Gra | 4.64 | **0.0000** |
| CHRE - CTST | Obl - Gra | 4.37 | **0.0000** |
| CHCI - CHLU | Fac - Obl | -1.79 | 0.0646 |
| CHCI - CHOR | Fac - Obl | -1.66 | 0.0762 |
| CHCI - CHRE | Fac - Obl | -2.21 | **0.0311** |
| CHPE - CHRE | Fac - Obl | -1.68 | 0.0777 |
| CHCI - CHPE | Fac - Fac | -0.63 | 0.3296 |
| CHCI - CHSP | Fac - Fac | 1.49 | 0.1021 |
| CHPE - CHSP | Fac - Fac | 2.29 | **0.0265** |
| CHCI - WAT | Fac - Env | 2.73 | **0.0088** |
| CHCI - SED | Fac - Env | 2.06 | **0.0388** |
| CHPE - WAT | Fac - Env | 3.74 | **0.0004** |
| CHPE - SED | Fac - Env | 3.00 | **0.0041** |
| CHSP - WAT | Fac - Env | 1.23 | 0.1543 |
| CHSP - SED | Fac - Env | 0.50 | 0.3619 |
| CHCI - CTFL | Fac - Gra | 1.91 | 0.0528 |
| CHCI - CTST | Fac - Gra | 1.90 | 0.0524 |
| CHPE - CTFL | Fac - Gra | 2.75 | **0.0086** |
| CHPE - CTST | Fac - Gra | 2.66 | **0.0101** |
| CHSP - CTFL | Fac - Gra | 0.46 | 0.3691 |
| CHSP - CTST | Fac - Gra | 0.55 | 0.3570 |
| WAT - SED | Env - Env | -0.82 | 0.2772 |
| CTFL - WAT | Gra - Env | 0.72 | 0.3000 |
| CTFL - SED | Gra - Env | -0.01 | 0.4968 |
| CTST - WAT | Gra - Env | 0.53 | 0.3557 |
| CTST - SED | Gra - Env | -0.14 | 0.4625 |
| CTFL - CTST | Gra - Gra | 0.12 | 0.4608 |

**Table S4:**
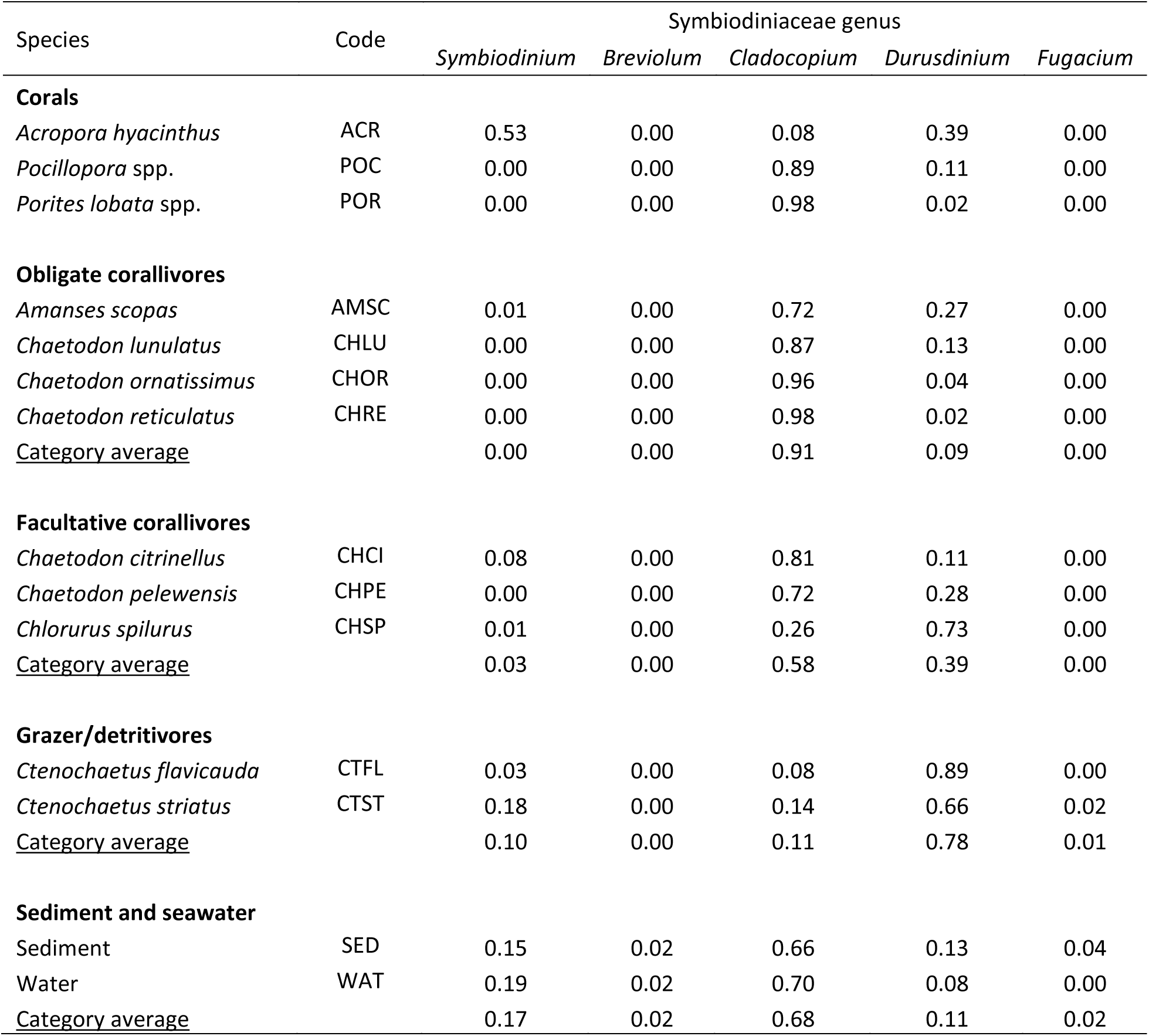
Mean relative abundances of genes associated with five Symbiodiniaceae genera identified in this study per sample type (e.g., *Amanses scopas* feces) and overall sample category (e.g., obligate corallivores) (Figure 3). Included samples (and sample sizes) for each sample category are as follows: Corals: *Acropora hyacinthus*, ACR (11); *Pocillopora* spp. = *Pocillopora* species complex, POC (12); *Porites lobata* spp. = *Porites lobata* species complex, POR (12). Obligate corallivores: *Amanses scopas*, AMSC (7)*; Chaetodon lunulatus*, CHLU (8); *Chaetodon ornatissimus*, CHOR (14); *Chaetodon reticulatus*, CHRE (11). Facultative corallivores: *Chaetodon pelewensis*, CHPE (8); *Chaetodon citrinellus*, CHCI (6); and *Chlorurus spilurus* CHSP (8). Grazer/Detritivores: *Ctenochaetus flavicauda*, CTFL (6); and *Ctenochaetus striatus*, CTST (6). Sediment and water: Sediment, SED (12); Water, WAT (7).

| Species | Code | Symbiodiniaceae genus |  |  |  |  |
| --- | --- | --- | --- | --- | --- | --- |
|  |  | *Symbiodinium* | *Breviolum* | *Cladocopium* | *Durusdinium* | *Fugacium* |
| Corals |  |  |  |  |  |  |
| *Acropora hyacinthus* | ACR | 0.53 | 0.00 | 0.08 | 0.39 | 0.00 |
| *Pocillopora* spp. | POC | 0.00 | 0.00 | 0.89 | 0.11 | 0.00 |
| *Porites lobata* spp. | POR | 0.00 | 0.00 | 0.98 | 0.02 | 0.00 |
| Obligate corallivores |  |  |  |  |  |  |
| *Amaneses scopas* | AMSC | 0.01 | 0.00 | 0.72 | 0.27 | 0.00 |
| *Chaetodon lunulatus* | CHLU | 0.00 | 0.00 | 0.87 | 0.13 | 0.00 |
| *Chaetodon ornatissimus* | CHOR | 0.00 | 0.00 | 0.96 | 0.04 | 0.00 |
| *Chaetodon reticulatus* | CHRE | 0.00 | 0.00 | 0.98 | 0.02 | 0.00 |
| <u>Category average</u> |  | 0.00 | 0.00 | 0.91 | 0.09 | 0.00 |
| Facultative corallivores |  |  |  |  |  |  |
| *Chaetodon citrinellus* | CHCI | 0.08 | 0.00 | 0.81 | 0.11 | 0.00 |
| *Chaetodon pelewensis* | CHPE | 0.00 | 0.00 | 0.72 | 0.28 | 0.00 |
| *Chlorurus spilurus* | CHSP | 0.01 | 0.00 | 0.26 | 0.73 | 0.00 |
| <u>Category average</u> |  | 0.03 | 0.00 | 0.58 | 0.39 | 0.00 |
| Grazer/detritivores |  |  |  |  |  |  |
| *Ctenochaetus flavicauda* | CTFL | 0.03 | 0.00 | 0.08 | 0.89 | 0.00 |
| *Ctenochaetus striatus* | CTST | 0.18 | 0.00 | 0.14 | 0.66 | 0.02 |
| <u>Category average</u> |  | 0.10 | 0.00 | 0.11 | 0.78 | 0.01 |
| Sediment and seawater |  |  |  |  |  |  |
| Sediment | SED | 0.15 | 0.02 | 0.66 | 0.13 | 0.04 |
| Water | WAT | 0.19 | 0.02 | 0.70 | 0.08 | 0.00 |
| <u>Category average</u> |  | 0.17 | 0.02 | 0.68 | 0.11 | 0.02 |

**Table S5:** Results from pairwise PERMANOVA tests on Symbiodiniaceae community composition at the genus level, based on Bray-Curtis distances (Figure 3). Samples were randomly subsampled from each sample category (n=12 each, see **Supplementary Materials** for sample names). We controlled for false positives with the Benjamini-Hochberg procedure. Significant p-values (<0.05) are bolded. Overall test results: df=6, F=17.3, R^2^=0.58, p=0.001 (PERMANOVA). *Acropora*: *Acropora hyacinthus*; *Pocillopora* spp. *= Pocillopora* species complex; *Porites lobata* spp. = *Porites lobata* species complex.

| Comparisons | R<sup>2</sup> | Adjusted p-value |
| --- | --- | --- |
| Obligate corallivore vs Facultative corallivore | 0.20 | **0.025** |
| Obligate corallivore vs Grazer/detritivore | 0.80 | **0.002** |
| Obligate corallivore vs Sediment and water | 0.14 | **0.020** |
| Obligate corallivore vs *Acropora hyacinthus* | 0.49 | **0.002** |
| Obligate corallivore vs *Pocillopora* spp. | 0.03 | 0.431 |
| Obligate corallivore vs *Porites lobata* spp. | 0.11 | 0.129 |
| Facultative corallivore vs Grazer/detritivore | 0.50 | **0.002** |
| Facultative corallivore vs Sediment and water | 0.09 | 0.117 |
| Facultative corallivore vs *Acropora hyacinthus* | 0.32 | **0.002** |
| Facultative corallivore vs *Pocillopora* spp. | 0.16 | **0.049** |
| Facultative corallivore vs *Porites lobata* spp. | 0.39 | **0.002** |
| Grazer/detritivore vs Sediment and water | 0.51 | **0.002** |
| Grazer/detritivore vs *Acropora hyacinthus* | 0.26 | **0.014** |
| Grazer/detritivore vs *Pocillopora* spp. | 0.82 | **0.002** |
| Grazer/detritivore vs *Porites lobata* spp. | 0.90 | **0.002** |
| Sediment and water vs *Acropora hyacinthus* | 0.28 | **0.002** |
| Sediment and water vs *Pocillopora* spp. | 0.14 | **0.006** |
| Sediment and water vs *Porites lobata* spp. | 0.25 | **0.002** |
| *Acropora hyacinthus* vs *Pocillopora* spp. | 0.49 | **0.002** |
| *Acropora hyacinthus* vs *Porites lobata* spp. | 0.57 | **0.002** |
| *Pocillopora* spp. vs *Porites lobata* spp. | 0.33 | **0.009** |

**Table S6:**
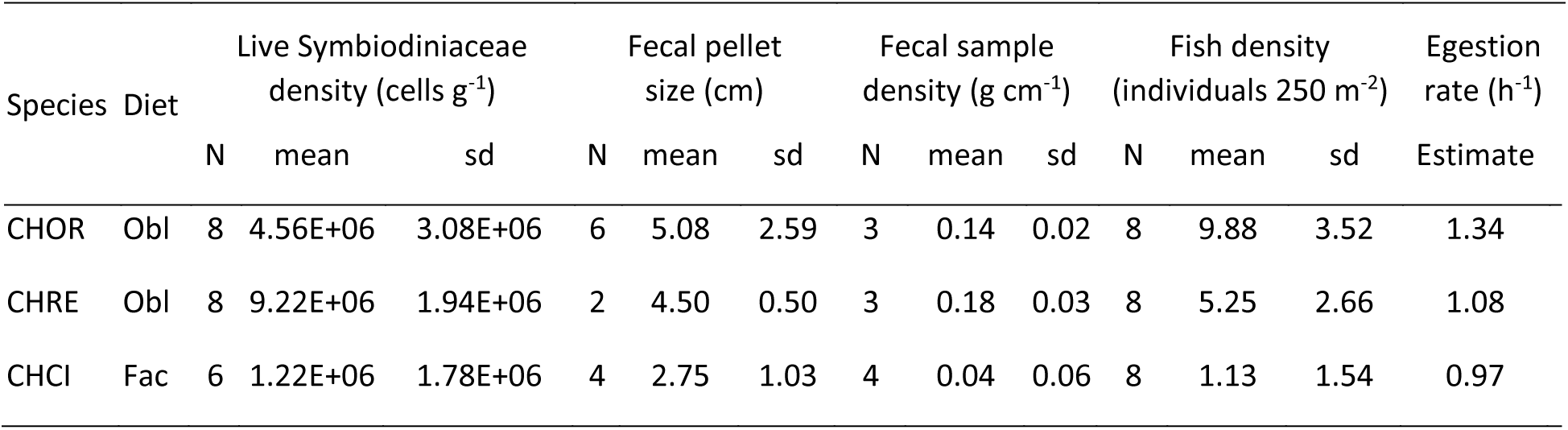
Overview of values used in bootstrap estimate of reef-scale Symbiodiniaceae dispersal (Figure 4). Fecal samples were used to calculate mean densities of live Symbiodiniaceae cells and mean fecal densities. Fish densities were calculated from the MCR LTER dataset (http://mcrlter.msi.ucsb.edu/cgi-bin/showDataset.cgi?docid=knb-lter-mcr.6 accessed February 14, 2020). Data on fecal pellet sizes and egestion rates were collected during *in situ* fish follows. Obl: obligate corallivore; Fac: facultative corallivore. See **Supplementary Methods** for additional details.

| Species | Diet | Live Symbiodiniaceae density (cells g<sup>-1</sup>) |  |  | Fecal pellet size (cm) |  |  | Fecal sample density (g cm<sup>-1</sup>) |  |  | Fish density (individuals 250 m<sup>-2</sup>) |  |  | Egestion rate (h<sup>-1</sup>) |
| --- | --- | --- | --- | --- | --- | --- | --- | --- | --- | --- | --- | --- | --- | --- |
|  |  | N | mean | sd | N | mean | sd | N | mean | sd | N | mean | sd | Estimate |
| CHOR | Obl | 8 | 4.56E+06 | 3.08E+06 | 6 | 5.08 | 2.59 | 3 | 0.14 | 0.02 | 8 | 9.88 | 3.52 | 1.34 |
| CHRE | Obl | 8 | 9.22E+06 | 1.94E+06 | 2 | 4.50 | 0.50 | 3 | 0.18 | 0.03 | 8 | 5.25 | 2.66 | 1.08 |
| CHCI | Fac | 6 | 1.22E+06 | 1.78E+06 | 4 | 2.75 | 1.03 | 4 | 0.04 | 0.06 | 8 | 1.13 | 1.54 | 0.97 |

